# CRISPR with independent transgenes is a safe and robust alternative to autonomous gene drives in basic research

**DOI:** 10.1101/017384

**Authors:** Fillip Port, Nadine Muschalik, Simon L Bullock

## Abstract

CRISPR/Cas technology allows rapid, site-specific genome modification in a wide variety of organisms. CRISPR components produced by integrated transgenes have been shown to mutagenise some genomic target sites in *Drosophila melanogaster* with high efficiency, but whether this is a general feature of this system remains unknown. Here, we systematically evaluate available CRISPR/Cas reagents and experimental designs in *Drosophila*. Our findings allow evidence-based choices of Cas9 sources and strategies for generating knock-in alleles. We perform gene editing at a large number of target sites using a highly active Cas9 line and a collection of transgenic gRNA strains. The vast majority of target sites can be mutated with remarkable efficiency using these tools. We contrast our method to recently developed autonomous gene drive technology for genome engineering (Gantz & Bier, 2015) and conclude that optimised CRISPR with independent transgenes is as efficient, more versatile and does not represent a biosafety risk.

## Introduction

The Clustered Regularly Interspaced Short Palindromic Repeat/CRISPR associated (CRISPR/Cas) genome engineering system is currently revolutionising experimental and applied biology (Doudna and Charpentier, 2014; Hsu et al., 2014). The system consists of the bacterial endonuclease Cas9 and a small chimeric guide RNA (gRNA), which directs Cas9 to a genomic target site 5’ to an NGG ‘protospacer adjacent motif’ (PAM) (Jinek et al., 2012). Repair of CRISPR/Cas-mediated DNA double strand breaks (DSBs) by endogenous machinery can result in mutagenic insertions and deletions (Indels) or – together with a supplied piece of donor DNA – precise sequence alterations. This methodology has been used successfully to edit the genome of a variety of organisms, although mutagenesis efficiencies have varied widely between different target sites, even in individual studies.

It has been proposed that inserting a cassette encoding Cas9 and gRNA into the endogenous target site of the gRNA will efficiently autocatalyse its integration into the homologous allele, thus converting heterozygous into homozygous cells (Esvelt et al., 2014; Oye et al., 2014). Such CRISPR/Cas ‘gene drive’ systems could potentially be used to efficiently spread traits within wild populations of plants and animals to address global problems in public health, sustainable agriculture, and environmental management (Burt, 2003; Esvelt et al., 2014; Sinkins and Gould, 2006). Gantz and Bier recently demonstrated such a ‘gene drive’ system using *Drosophila melanogaster*, a major model organism in biomedical research (Gantz and Bier, 2015). They propose that by initiating a ‘mutagenic chain reaction’ (MCR) their technology can be used to accelerate laboratory genome engineering and reveal homozygous mutant phenotypes in genetic screens. However, there are major biosafety concerns about the use of autonomous gene drive technology in the laboratory because of the risk of accidental infiltration of autocatalytic alleles into wild populations (Esvelt et al., 2014; Oye et al., 2014).

Here we describe the systematic evaluation of *Drosophila* CRISPR/Cas tools and experimental designs that do not create gene drives. Our work reveals that optimised CRISPR with independent, integrated Cas9 and gRNA transgenes can achieve remarkably efficient germ line and somatic gene targeting at a very large proportion of genomic target sites. The performance of these methods is comparable to that reported for Gantz and Bier’s MCR technology. We conclude that transgenic CRISPR/Cas is a safe and consistently efficient method to modify the fly genome and encourage others to apply similar technology to other experimental model organisms.

## Results

We and others have shown that transgenic *cas9* strains can in principle be used to rapidly and efficiently modify the *Drosophila* genome in somatic and germ line cells (Chen et al., 2015; Gratz et al., 2014; Kondo and Ueda, 2013; Port et al., 2014; Ren et al., 2013, 2014; Sebo et al., 2013; Xue et al., 2014; Zhang et al., 2014). gRNAs can be delivered into *cas9* strains by injection of plasmids or mRNA, or can be encoded by a previously integrated transgene (which we now refer to as CRISPR with independent transgenes (CRISPR-it)). There is significant uncertainty in the fly community about which of the many published *cas9* strains are most effective for genome engineering experiments.

We evaluated the performance of all transgenic *cas9* lines available at the time of study. The collection consisted of six lines that we assessed previously (Port et al., 2014) and eight others (Suppl. Table 1). The transgenes differed in regulatory elements, *cas9* codon usage and genomic integration sites (Suppl. Table 1). We crossed *cas9* flies to a previously validated, ubiquitously expressed gRNA transgene (*U6:3-gRNA-e)*, which targets the 5’ end of the coding sequence (CDS) of the pigmentation gene *ebony* (*e*) (Port et al., 2014). Phenotypic assays using this gRNA typically report only on out-of-frame Indels, as most in-frame mutations at the *gRNA-e* target site retain protein function (Port et al., 2014). All adult progeny expressing *U6:3-gRNA-e* and *cas9* under the control of either the *actin5c* (*act*) or *vasa* promoter had a large proportion of cuticle that was dark, demonstrating efficient biallelic disruption of *e* in somatic cells (Figure 1A). In contrast, seven of the nine lines tested expressing *cas9* under *nanos* (*nos*) regulatory elements did not have detectable somatic activity when crossed to *U6:3-gRNA-e* (Figure 1A). These findings confirm and extend our previous findings on the expression patterns of *cas9* driven by these classes of regulatory elements (Port et al., 2014). Phenotypic analysis of the progeny of *cas9 U6:3-gRNA-e* flies and *e* mutant partners revealed a wide range of germ line transmission rates of CRISPR/Cas-induced non-functional *e* alleles, varying between 0.5% and 49% (Figure 1B). *act-cas9*, *vasa-cas9* BL51323 and *nos-cas9* TH00787.N (which we refer to as TH_attP2) gave rise to similar, high rates of mutagenesis.

**Figure 1:**
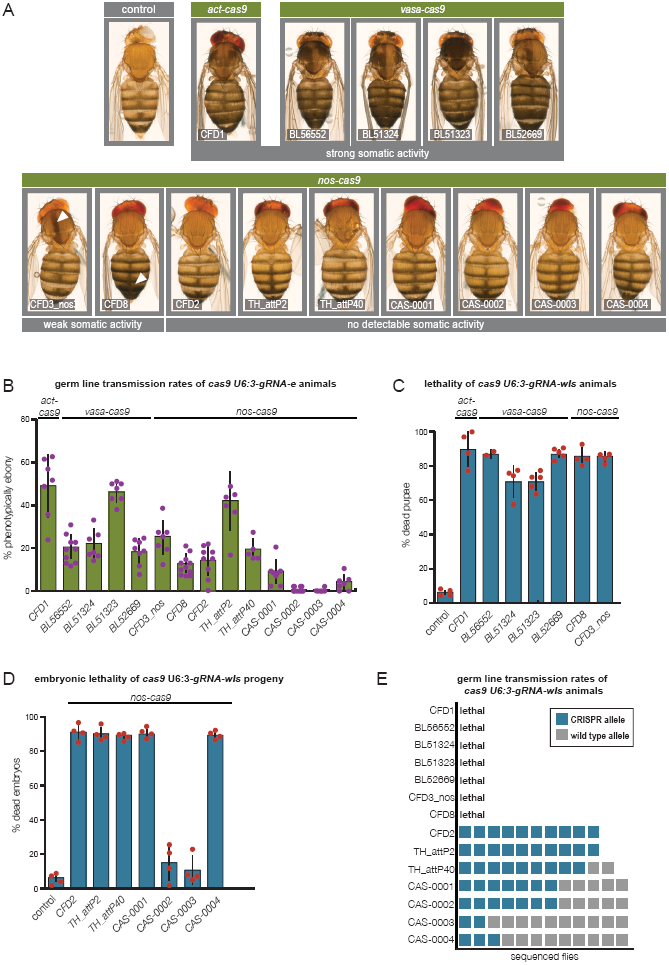
Large variation of somatic and germ line activity in publicly available transgenic *cas9* strains. (**A**) Representative examples of female flies expressing one copy of the different *cas9* transgenes and one copy of *U6:3-gRNA-e* transgene (> 100 adults of each genotype were examined). Darker body coloration indicates CRISPR/Cas-mediated mutagenesis of e in epidermal cells (arrowheads: sporadic biallelic targeting of e). Assessment of germ line transmission of non-functional e alleles from animals expressing different cas9 transgenes and the *U6:3-gRNA-e* transgene. In B – D, dots show data from individual crosses with mean ± standard deviation (SD) represented by the underlying bar chart. In B, typically between 30 and 140 progeny were analysed for each cross Frequency of pupal lethality caused by *cas9* transgenes with somatic activity in combination with the *U6:3-gRNA-wls* transgene. Flies that eclosed from the pupal case had leg and wing defects consistent with reduced *wls* function and died shortly afterwards. In C and D, control flies were of the genotype *act-cas9 U6:3-gRNA-e*, which have widespread induction of DSBs. Typically, 50 - 150 animals were examined for each cross. (**D**) Percentage of non-viable embryos laid by females expressing different germ line restricted *nos-cas9* transgenes and the *U6:3-gRNA-wls* transgene following mating to wild-type males. Typically, between 100 and 300 embryos were examined for each cross. (**E**) Assessment of germ line transmission rates of CRISPR/Cas-induced *wls* mutations. Offspring from *nos-cas9 U6:3-gRNA-wls* male flies crossed to wild-type females were genotyped by PCR of the *wls* locus followed by sequencing. Each square represents an individual genotyped fly. Many mutations were out-of-frame, presumably disrupting wls function. For the *nos-cas9* CAS series of lines there was little correlation between the rate of embryonic lethality induced by *U6:3-gRNA-wls* and the rate of germ line transmission of *wls* alleles to the viable offspring. Germ line transmission depends on a mutation of *wls* in the transcriptionally quiescent oocyte nucleus, whereas embryonic lethality arises from mutagenesis of the gene in the auxiliary nurse cells, which supply protein products to the egg through cytoplasmic bridges. These *nos-cas9* lines may therefore be differentially active in targeting *wls* in the nurse cells and oocyte within the female germ line.

We also evaluated the *cas9* lines with a second gRNA transgene (*U6:3-gRNA-wls*) targeting the essential gene *wntless* (*wls*) (Port et al., 2014). As expected, *cas9* transgenic strains that had somatic activity in combination with *U6:3-gRNA-e* resulted in non-viable progeny when crossed to *U6:3-gRNA-wls* (Figure 1C). The progeny of the other seven *nos-cas9* lines were viable in combination with *U6:3-gRNA-wls*, although a small proportion of adults from these crosses had wing defects indicative of gene targeting in a small subset of somatic cells (Figure 1; Suppl. Figure 1). Targeting of *wls* in the germ line was assessed by crossing *nos-cas9 U6:3-gRNA-wls* females to wild-type males. All but two of the *nos-cas9* transgenes resulted in a high proportion of embryos arresting with defective segmentation (Figure 1D), a phenotype associated with biallelic disruption of *wls* in the female germ line (Bänziger et al., 2006; Bartscherer et al., 2006). All crosses gave rise to some viable offspring, with genotyping of the *wls* locus from these flies revealing substantial variation in the frequency of germ line transmission of CRISPR/Cas-induced mutations (Figure 1C). In two cases, all analysed offspring (10/10) received a modified *wls* allele. Together these results reveal that the *cas9* lines differ substantially in their somatic and germ line activity. Whereas *act-cas9* and *vasa-cas9* lines can directly reveal null mutant phenotypes in the soma when combined with gRNA transgenes, a subset of *nos-cas9* lines (e.g. CFD2, TH00788.N (referred to as TH_attP40) and TH_attP2 (inserted on the X, 2^nd^ and 3^rd^ chromosomes, respectively)) can efficiently generate Indel mutations in both essential and non-essential genes in the germ line.

Another important application of CRISPR/Cas is the precise modification of the genome by homology directed repair (HDR). This involves Cas9-mediated induction of DSBs at the target site in the presence of exogenous donor DNA. Three strategies are particularly appealing in *Drosophila* because of their relative simplicity. Embryos that are transgenic for both *cas9* and *gRNA* can be injected with donor DNA (Port et al., 2014), transgenic *cas9* embryos can be injected with a mixture of donor DNA and gRNA-encoding plasmid (Gratz et al., 2014; Ren et al., 2014; Zhang et al., 2014) or non-transgenic embryos can be injected with a donor plasmid together with a plasmid encoding Cas9 and gRNA (Gokcezade et al., 2014). No study has directly compared the efficiency with which all three methods facilitate HDR with the same gRNA and donor construct. To do so we constructed a donor plasmid designed to knock-in an expression cassette containing red fluorescent protein (RFP) under an eye specific promoter into the essential *wg* gene (Figure 2 – Suppl. Figure 2). The plasmid, which contained *wg* homology arms of 1.4 and 1.7 kb, was used in circular form in combination with a previously validated gRNA (*gRNA-wg* (Port et al., 2014)). For technical reasons (see Figure 2 legend) we only followed integration of donor DNA using male flies.

**Figure 2:**
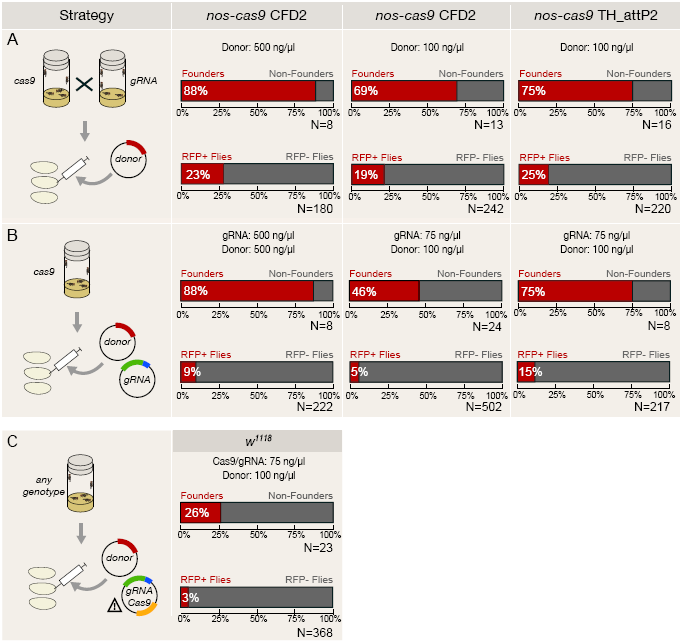
Assessing strategies for generating knock-in alleles with CRISPR/Cas-mediated HDR. (**A**) *nos-cas9* female flies of the indicated strain were crossed to *U6:3-gRNA-wg* males and a donor plasmid encoding RFP was injected into the embryonic progeny at the indicated concentrations. Due to a pre-existing X-linked RFP insertion in the CFD2 stock only male G_0_ and F_1_ flies could be analysed. For consistency we only analysed male flies in all other HDR experiments. G_0_ males that give rise to RFP positive offspring were designated ‘founders’. The percentage of male progeny with RFP expression relative to all offspring (i.e. from fertile founder and non-founder G_0_ males) is indicated below. N, total number of males analysed. (**B**) Donor and *U6:3-gRNA-wg* plasmids were injected into *nos-cas9* embryos. In B and C, plasmids were mixed to give an injection mix containing the concentrations shown. (**C**) Non-transgenic *w*^*1118*^ embryos were injected with donor DNA and a single plasmid containing both *hsp70-cas9* and *U6:2-gRNA-wg*. The attention sign indicates that unintended integration of this plasmid at the gRNA target site could create an autonomous gene drive. Propagation of a gene drive is not possible in our experiment as integration at the *gRNA-wg* target site would create a lethal allele. Suppl. Table 2 contains detailed results for all HDR experiments.

For the experiments involving transgenic *cas9* supply we used different plasmid concentrations and two different *nos-cas9* strains (CFD2 and TH_attP2) (Figure 2). Injection of the donor plasmid into *cas9 gRNA-wg* double transgenic embryos resulted in 19 – 25% of all offspring from G_0_ flies having integration of the RFP construct, compared to 5 – 15% when the *U6:3-gRNA-wg* plasmid was injected with donor DNA into *nos-cas9* embryos (Figure 2A, B and Suppl. Table 2).

We assessed the efficiency of plasmid-based delivery of Cas9 and gRNA using a published vector that encodes both components (Gokcezade et al., 2014). Although expected to be a rare event, accidental integration of such a plasmid at the gRNA target site would create an autonomous gene drive. We were able to safely test this plasmid as insertion at the *gRNA-wg* target site would be homozygous lethal, preventing the allele from being propagated. Upon co-injection of the *cas9*/*gRNA* plasmid and donor plasmid into non-transgenic embryos, only 3% of all progeny from G_0_ flies had genomic integration of the RFP cassette (Figure 2C and Suppl. Table 2). Integration of the donor DNA within the *wg* locus could be confirmed for 87 of 98 RFP positive flies tested from our set of HDR experiments (Figure 2 – Suppl. Figure 2 and Suppl. Table 2). In summary, the methods using a transgenic *cas9* source resulted in the highest frequency of knock-in alleles, with injection into embryos that are also transgenic for the gRNA giving the most efficient targeting.

The above results, together with previous literature (Kondo and Ueda, 2013; Port et al., 2014), provides evidence that transgenic supply of both Cas9 and gRNA can lead to very high rates of mutagenesis. An important outstanding question is whether the high rates of efficiency observed for a small number of genomic target sites in these studies are a general feature of this system. Previous studies have found that gRNAs delivered by injection of RNA or a plasmid can differ substantially in their activity (Bassett et al., 2013; Lee et al., 2014; Ren et al., 2014; Zhang et al., 2014), prompting some to recommend screening for active gRNAs in cell culture before performing whole animal experiments (Zhang et al., 2014).

**Figure 3:**
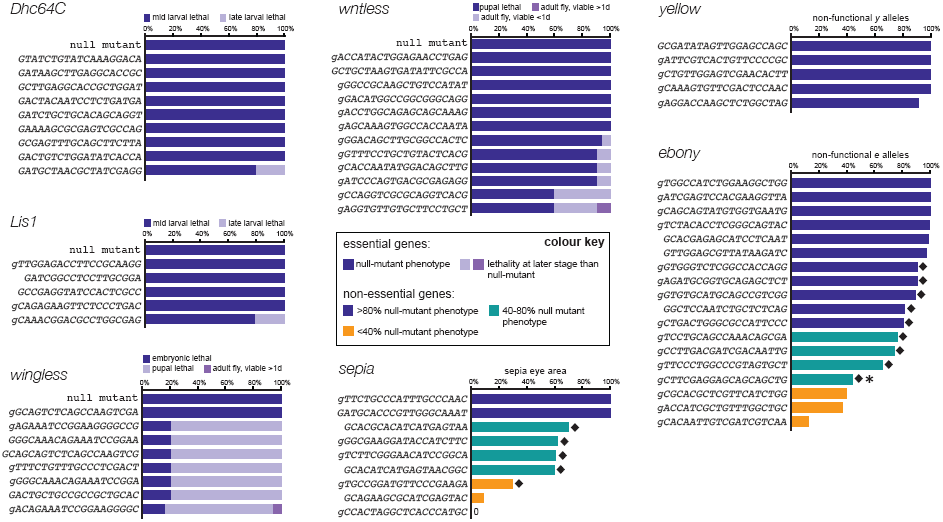
Almost all transgenic gRNAs are highly active with a strong transgenic source of Cas9. Male flies transgenic for individual gRNAs were crossed to *act-cas9* females. Somatic phenotypes of act-cas9 gRNA progeny are summarised for gRNAs targeting *Dhc64C*, *Lis1*, *wg*, *wls* and *se*. The *gRNA-se* with no detectable activity was found to contain a polymorphism. Activity in the germ line is shown for gRNAs targeting *y* and *e* (data show mean percentage of progeny that inherited a non-functional allele from three independent crosses (see Suppl. Table 3 for more details and somatic phenotypes of these gRNAs)). gRNAs highlighted by black diamonds frequently give rise to functional in-frame mutations (Suppl. Table 3), suggesting that their efficiency in creating Indel mutations often approaches 100%. The gRNA additionally highlighted by an asterisk gives rise to an unexpected large number of in-frame mutations in functional alleles (8/8 flies analysed), suggesting a micro-homology mediated bias in non-homologous end joining. Since transcription initiation from U6 promoter is proposed to require a G nucleotide, we extended gRNAs by a mismatched G where necessary (lower case). gRNAs with spacer sequences of 18 nt, 19 nt and 20 nt can support efficient mutagenesis, consistent with previous evidence that short truncations of spacers usually do not strongly affect activity of gRNAs (Fu et al., 2014; Ren et al., 2014).

To determine what fraction of gRNAs are functional in the CRISPR-it system we generated a set of 66 transgenic fly strains each expressing a different gRNA. The gRNAs were designed to target seven genes including essential genes (*wingless* (*wg*), *wls*, *Lissencephaly-1* (*Lis1*), *Dynein heavy chain at 64C* (*Dhc64C*)) and non-essential pigmentation genes (*e*, *yellow* (*y*; also controlling coloration of the cuticle), and *sepia* (*se*; controlling coloration of the eye)). gRNA target sites were based on the reference sequence of the *Drosophila* genome (Flybase release 6.02). They were located in the 5’ half of the coding sequence of each gene, such that out-of-frame Indels are likely to create loss-of-function alleles, but were otherwise selected at random. gRNAs were expressed under the control of the strong, ubiquitous *U6:3* promoter (Port et al., 2014) and integrated at the same genomic location to ensure comparable expression of all transgenes.

The 66 individual gRNA strains were crossed to the *act-cas9* line, which has ubiquitous, high activity (Figure 1A,B). Strikingly, analysis of the progeny revealed that all but one of the gRNAs gave rise to a somatic mutant phenotype (Figure 3 and Suppl. Table 3). Sequencing of our strains revealed a single nucleotide polymorphism in the target site of the inactive gRNA, designed to target *se*, providing a likely explanation for its inactivity. Most of the gRNAs targeting the essential genes caused lethality at the same developmental stage as reported for the respective homozygous null-mutant animals, with the remainder leading to developmental arrest at later stages (Figure 3). With the exception of the single inactive *gRNA-se* transgene, all of the gRNAs targeting non-essential genes gave adults with regions of tissue with the null-mutant pigmentation phenotype (Figure 3, Figure 3 – Suppl. Figure 3 and Suppl. Table 3). For almost all gRNAs, biallelic gene disruption was usually observed in over 50% of the affected tissue, with half of the gRNAs, including all five targeting *y*, leading to more than 80% mutant tissue (Suppl. Table 3). Non-specific phenotypes were not observed in *act-cas9* adults co-expressing any of the gRNA transgenes targeting *y*, *e* or *se*, consistent with previous evidence that CRISPR/Cas operates with substantial fidelity in *Drosophila* (Bassett et al., 2013; Gratz et al., 2014; Ren et al., 2014). The above results demonstrate that the vast majority of randomly selected gRNAs efficiently induce biallelic loss-of-function mutations in target genes, thereby directly revealing the mutant phenotype. There was, however, variation in the strength of the somatic phenotypes observed for gRNAs targeting the same gene (Figure 3 and Suppl. Table 3). Weaker phenotypes could arise either through a reduced frequency of induced Indel mutations or because functional in-frame mutations at the specific gRNA target site provide a significant fraction of cells with functional protein.

To directly test the hypothesis that in-frame mutations can mask high rates of Indel induction by some gRNAs, we characterised CRISPR/Cas-induced mutations transmitted to the next generation. Flies expressing *act-cas9* and a transgenic gRNA targeting *y* or *e* were crossed to *y* or *e* mutant flies, respectively (Figure 3). The rates of transmission of non-functional alleles across target sites was generally higher than reported when gRNAs were delivered by plasmid injection into transgenic *cas9* embryos (Ren et al., 2014) (Figure 3 – Suppl. Figure 4A,B), providing further evidence of the relatively high activity of CRISPR-it. All five *gRNA-y* and six of 18 *gRNA-e* transgenes transmitted non-functional alleles to >93% of their offspring (Figure 3). The high frequency of non-functional alleles strongly suggests that in-frame mutations at these target sites usually disrupt protein function. Transmission rates for non-functional mutations ranged from 13 – 90% for the other 12 *gRNA-e* lines (Figure 3). To explore the basis of this variation, we determined the sequence of *e* around the target sites in some of the phenotypically wild-type progeny from the 12 lines. For nine of the gRNAs, in-frame Indel mutations were detected in all, or almost all, analysed animals (Suppl. Table 3; Figure 3). Thus these gRNAs appear to create Indels in the vast majority of *e* alleles, with differences in the functional effects of in-frame mutations at their target sites influencing the outcome of phenotypic assays. We confirmed that this effect was not specific for target sites in *e* by molecular analysis of the *se* locus in offspring of *act-cas9 gRNA-se* flies, which again revealed in-frame mutations in the majority of functional alleles (Suppl. Table 3; Figure 3).

Indel mutations were not readily found in functional alleles inherited from flies expressing *act-cas9* and the three *gRNA-e* transgenes that were the least efficient in the phenotypic assays (transmission of non-functional alleles in 13 – 35% of cases) or one poorly active *se* gRNA (7% of phenotypically mutant eye tissue) (Suppl. Table 3; Figure 3). Thus, these gRNAs induce Indels with relatively low frequency. Sequencing of the target site for these gRNAs excluded the possibility that polymorphisms were responsible for their reduced activity. gRNAs with lower activity were not readily predicted by available online gRNA design tools and their target sites did not have a low GC content in the region proximal to the PAM (Figure 3 – Suppl. Figure 4C – J; Suppl. Table 4), a parameter which strongly predicted less active gRNAs encoded by plasmids injected into transgenic *cas9* embryos (Ren et al., 2014). Several gRNAs targeting sites with relatively low PAM proximal GC content had very high activity in our experiments (Figure 3 – Suppl. Figure 4A, F, J). Collectively, our results demonstrate that optimised CRISPR-it is a very robust genome engineering system, leading to highly effective mutagenesis at the vast majority of target sites.

## Discussion

We set out to provide a systematic evaluation of tools and design parameters for transgenic CRISPR/Cas genome engineering in *Drosophila melanogaster*. We show that selection of the *cas9* strain is critical for the successful application of this methodology, as publicly available lines vary widely in their activity in somatic and germ line cells. Success rates also vary substantially between different protocols for HDR-mediated knock-ins. High efficiency is particularly important for HDR without selectable markers, which introduce unwanted sequences into the target locus, as in most cases integration must be detected by PCR-based analysis of genomic DNA. Most importantly, we show that use of transgenic gRNAs allows highly efficient mutagenesis across a large proportion of target sites, with our analysis of germ line transmission of mutations in pigmentation genes suggesting that the vast majority of gRNAs cause close to maximal rates of Indel induction. Incomplete penetrance of phenotypes induced by some transgenic gRNAs is due to functional in-frame mutations; co-expression of more than one gRNA using a dual gRNA vector (Port et al., 2014) should alleviate this problem.

Importantly, our study highlights that CRISPR-it is a safe, efficient and robust alternative to the recently reported MCR, which uses an autonomous gene drive for functional studies (Gantz and Bier, 2015) (Table 1). MCR has so far only been demonstrated at a single target site in the *Drosophila y* gene, a locus often used in pioneer studies and which appears to have an unusually high susceptibility for gene targeting (Gratz et al., 2013; Rong and Golic, 2000). MCR can rapidly reveal recessive somatic phenotypes as heterozygous mutations are efficiently converted to the homozygous state. We show here that CRISPR-it can induce biallelic gene disruption with similar efficiency in the soma using many target sites, including five in *y* (Figure 3 and Suppl. Table 3). Because this method uses separate *cas9* and *gRNA* transgenes, mutagenesis is inactivated by breeding. This is not the case for the MCR system due to linkage of the *cas9* and *gRNA* sequences at the target locus. This means that escape of flies from the laboratory could conceivably result in rapid spread of an MCR allele in the wild, with unpredictable ecological consequences. Such a risk is in principle also associated with other plasmids encoding both Cas9 and gRNA (e.g. Gokcezade et al., 2014), although the risk of insertion at the gRNA target site is much lower due to the absence of homology arms.

**Table 1:** Comparison of MCR and CRISPR-it technology.

|  | Mutagenic chain reaction | CRISPR-it |
| --- | --- | --- |
| **Work environment** | Physical containment<br>• Triple contained flies<br>• Locked facilities at all times<br>• Accounting for individual flies<br>• Single experimenter | Regular fly room and working procedures |
| **CRISPR components** | On one construct | Separate, integrated transgenes |
| **Multiplexing<sup>a</sup>** | No | Yes |
| **Directly reveals recessive phenotypes** | Yes<br>(so far shown for a single target site) | Yes<br>(shown for many target sites) |
| **Efficiency of germ line transmission of mutations** | Up to 100% | Up to 100% |
| **CRISPR inactivated by crossing** | No | Yes |
| **Risk of infiltrating wild and lab populations** | Yes | No |
| **Genotype at target site** | Mosaic in all generations<br>(mostly homozygous for MCR allele) | F1: Mosaic<br>(mostly biallelic for Indels)<br>F2: heterozygous<br>F3: homozygous |
| **Time from cloning to recessive phenotype** | ~25 days | ~36 days<sup>b</sup> |
| **Targeting essential genes** | No<sup>c</sup> | Yes |
| **Tissue specific** | No | Yes |

a: Generation of multiple gRNA plasmids can be multiplexed during cloning and transgenesis. Multiplexing is not possible during MCR. b: One additional generation. c: Would require complex rescue construct (Gantz and Bier, 2015).

Working with flies containing an MCR cassette has to be performed using very strict biosafety procedures (Table 1) (Gantz and Bier, 2015). No such precautions are necessary for transgenic CRISPR work, meaning that experiments can be performed much more conveniently and rapidly. Furthermore, CRISPR-it can readily reveal mutant phenotypes of essential genes and can be employed in a tissue specific manner (Port et al., 2014; Xue et al., 2014), applications that are not easy to achieve with MCR. The ability of CRISPR-it to efficiently reveal recessive mutant phenotypes associated with a large proportion of genomic target sites also makes it a highly attractive system for large scale F_1_ mutagenesis screens.

Gene drive technology has great promise for applications such as ecosystem management and pest control and may well be adopted for basic research applications (Burt, 2003; Esvelt et al., 2014; Sinkins and Gould, 2006). However, responsible use of this technology will require development of robust molecular containment strategies (Esvelt et al., 2014). We therefore urge researchers not to apply gene drives without these safeguards in place. Our experiments illustrate safe and effective genome engineering strategies that are already available for basic research and can be applied to any genetically tractable organism.

## Methods

### Plasmid construction

All primer sequences (purchased from (Integrated DNA Technologies (IDT)) are listed in Suppl. Table 5. Unless stated otherwise, enzymatic reactions were performed according to the manufacturers’ guidelines. To generate gRNA expression vectors, *pCFD3* (Port et al., 2014) (Addgene 49410) or *pDCC6* (Gokcezade et al., 2014) (Addgene 59985) were cut with *BbsI* and dephosphorylated with alkaline phosphatase, followed by gel purification of the linear plasmid. To introduce the target specific spacer sequence, two oligonucleotides containing the spacer sequence and reverse complement spacer sequence, as well as appropriate overhangs, were mixed (1μl each oligo (100μM stock solutions)) together with 1μl 10x T4 Ligation Buffer (New England Biolabs (NEB)), 6.5μl dH_2_O and 0.5 μl T4 polynucleotide kinase (NEB). Phosphorylation and annealing of oligos was performed in a thermo cycler (30 min at 37°C, 5 min at 95°C, followed by ramping down to 25°C at 5°C/min). Annealed oligos were diluted 1:200 in dH_2_O and ligated into the linear expression plasmids with T4 DNA ligase (NEB), followed by transformation into chemically competent bacteria. A step-by-step protocol is available from www.crisprflydesign.org.

### *Drosophila* culture

Flies were maintained at 25°C and 50% humidity at a 12h light/dark cycle.

### Assessing activity of *cas9* lines

To assess *cas9* line performance in targeting *e*, virgin females from the various *cas9* lines were mated to *U6:3-gRNA-e* transgenic males. The resulting double transgenic offspring was examined for CRISPR/Cas-induced phenotypes, and virgins heterozygous for both *cas9* and *U6:3-gRNA-e* were mated to *e* mutant males (*w;;TM3/TM6b*). The number of progeny from these crosses with ebony pigmentation was recorded. At least five independent crosses were analysed for each *cas9* line.

To analyse the ability of the various *cas9* lines to mediate mutagenesis of *wntless* (*wls*), virgin *cas9* females were crossed to *U6:3-gRNA-wls* transgenic males. At least three independent crosses were performed for each genotype. In cases where the resulting *cas9 U6:3-gRNA-wls* animals had significant rates of pupal lethality the number of dead versus total pupae was recorded. Males from the remaining, viable *cas9 U6:3-gRNA-wls* genotypes were crossed to *y w hs-FLP;;MKRS/TM6b* virgins. To determine the rate of germ line transmission, genomic DNA was isolated from some of the offspring, using microLysis-Plus (Microzone) according to the manufacturer’s instructions, and used for PCR analysis. Primers *wls_geno_fwd* and *wls_geno_rev* were used to amplify the regions flanking the gRNA-*wls* target site. The genomic sequence from 10 to 12 flies from two independent crosses was analysed for each genotype. To monitor embryonic viability of the offspring, virgins heterozygous for both *cas9* and *U6:3-gRNA-wls* were crossed to wild-type males, and the cross was transferred to cages mounted on apple juice agar plates. All embryos from a 1 – 2 h egg collection were counted and kept at 25°C for 48 h to allow for embryonic development to be completed. After 48 h the number of embryos that hatched was recorded. Where inspected, arrested embryos had segmentation defects. This analysis was carried out in quadruplicate for each genotype. Intercrosses of *act-cas9 U6:3-gRNA-e* flies were used as a control.

### Embryo injections and transgenesis

Microinjection into embryos were performed using standard procedures as described previously (Port et al., 2014). To generate transgenic gRNA lines, plasmids were often injected in pools (Bischof et al., 2013). Equal amounts of gRNA plasmids (5 – 20 per pool) were mixed and diluted to a final concentration of 150 ng/μl in dH_2_O and injected into *y v nos-PhiC31; attP40* (Bloomington Stock (BL)25709) embryos. Single transgenic offspring, selected by the *v*^+^ marker encoded by the gRNA plasmid, were mated with *v; Sp/CyO* flies for several days and then squashed in 10 μl microLysis-Plus (Microzone) to extract genomic DNA. To identify the integrated gRNA plasmid the transgene was amplified by PCR using primers *U63fwd1* and *CFD4seqrev3* and the resulting PCR product was submitted for Sanger sequencing (Source Bioscience) using the former primer.

### Assessing activity of the 66 gRNA transgenes

Transgenic gRNA males were crossed to *act-cas9* virgin females. Crosses with gRNAs targeting essential genes were monitored daily, followed by estimation of the proportion of dead offspring at each developmental stage. The activity of gRNAs targeting non-essential genes was monitored in adult offspring of *cas9* × *gRNA* crosses (note that we did not notice significant lethality before adulthood in any of these crosses). Flies expressing *act-cas9* and gRNAs targeting *se* were analysed one week after eclosion, when the difference in eye pigmentation between wild-type and *se* mutant tissue is most obvious (Figure 3 – Suppl. Figure 3). The percentage of eye tissue that was *se* mutant was analysed in at least 20 *act-cas9 gRNA-se* flies by three researchers blind to the genotype. The results were highly similar for each researcher and the average of the three independent recordings is presented in Figure 3. Flies expressing gRNAs targeting *y* or *e* and *act-cas9* were visually inspected for their respective pigmentation phenotype between three and six days after eclosion. Male flies were selected independently of the severity of the somatic pigmentation phenotype and crossed to either *y* or *e* mutant partners. The number of progeny with yellow or ebony pigmentation was recorded. Data presented in Figure 3 is the average of three independent crosses. The percentage of yellow offspring was normalised to account for the 50% of flies that were expected to be yellow due to the parental genotype.

### Image acquisition and processing

Flies were anaesthetized with CO_2_ and submerged in 90% ethanol/10% glycerol for at least 4 h, followed by mounting on Sylgard plates (Dow Corning) in 50% ethanol/50% glycerol. Images were captured with a Canon 550D camera equipped with a Canon 24 mm f1.8 lens mounted on a stereomicroscope (Leica MZFLIII). Camera settings were entirely manual and constant illumination was used in each session. Images presented in the same figure were captured in a single session. Brightness and contrast was adjusted with Adobe Photoshop software, with identical manipulations for each image within a series.

### Sequence analysis of CRISPR/Cas-induced mutations

Genomic DNA from single flies was extracted with 10μl microLysis-Plus (Microzone). 0.75 μl DNA solution was used as a template in 25 μl PCR reactions using the Q5 Hot Start 2x master mix (NEB). The sequence of the primers for the individual target genes is listed in Suppl. Table 5. PCR products were purified using the Qiagen PCR purification kit and submitted to Sanger sequencing using the forward PCR primer. Sequencing chromatograms from flies that inherited a CRISPR/Cas-induced Indel and are consequently heterozygous at the gRNA target site contained an overlay of chromatograms from both alleles. Chromatograms were analysed with Tide (http://tide.nki.nl/; (Brinkman et al., 2014)) and re-examined manually to correct for possible software mistakes, which occur occasionally for alleles involving insertions.

### Assessing different HDR strategies

Primers *pBSwg3’HA_fwd* and *pBS-wg5’HA_rev* were used to amplify the 5’ and 3’ *wg* homology arms and the pBluescript SK-(+) vector backbone from the previously described *Wg::GFP* donor plasmid (Port et al., 2014). The 3xP3-RFP-!tub-3’UTR (gift from Nick Lowe and Daniel St Johnston) sequence was amplified by PCR using primers *3xP3-RFP_fwd* and *3xP3-RFP_rev*. Both fragments were then joined by Gibson assembly to generate the RFP donor construct. Sequence-verified plasmid DNA was purified using the MinElute PCR purification kit from Qiagen. To test HDR efficiency using transgenic *cas9* and gRNA, the purified circular donor plasmid was injected into embryos resulting from crosses between *nos-cas9* CFD2 or *nos-cas9* TH_attP2 virgin females and *U6:3-gRNA-wg* males. For the protocol based on transgenic *cas9* stocks, both the donor and *U6:3-gRNA-wg* plasmid were injected into embryos either hemizygous (CFD2 males) or homozygous (CFD2 females or TH_attP2 males and females) for the *nos-cas9* transgene. To test HDR efficiency using the *pDCC6* plasmid encoding *cas9* and *U6:2-gRNA-wg* ((Gokcezade et al., 2014); hereafter called *pDCC6-wg)*, both donor and *pDCC6-wg* were injected into embryos of an isogenised *w*^*1118*^ stock (Bloomington Stock Centre ((BL)5905)). The concentrations of injected plasmids for the HDR experiments are given in Figure 2. Injected embryos were kept at 18°C for ∼ 48 h and then moved to 25°C. G_0_ males were crossed to *yw hs-FLP; Sp/CyO* females and the resulting male offspring was screened for the presence of RFP. To test whether the 3xP3-RFP cassette was inserted into the *wg* locus, genomic DNA was isolated from some of the RFP positive male offspring for PCR analysis as described above, with primer pairs *wgHRgeno_fwd1/wgHRgeno_rev1* and *wgHRgeno_fwd2*/*wgHRgeno_rev2* used to amplify the regions flanking each homology arm

## Acknowledgements

We would like to thank Sean Munro for encouragement and financial support of N.M., Peter Duchek, Nick Lowe and Daniel St Johnston for materials, Kevin Esvelt for discussions on gene drives, Neil Grant for help with photography, Mohammad Mofatteh for help with scoring eye phenotypes, users of www.crisprflydesign.org for feedback, and the *Drosophila* CRISPR community for freely sharing stocks and reagents. This study was supported by a Marie-Curie IntraEuropean Fellowship (to F.P.), a Wellcome Trust Strategic Award (095927/B/11/A) (N.M.) and a UK Medical Research Council Project U105178790 (to S.B.).

-Preprint design based on template by Denis Eckmeier www.eckmeier.de/ -

## Supplementary material

**Supplementary Figure 1:**
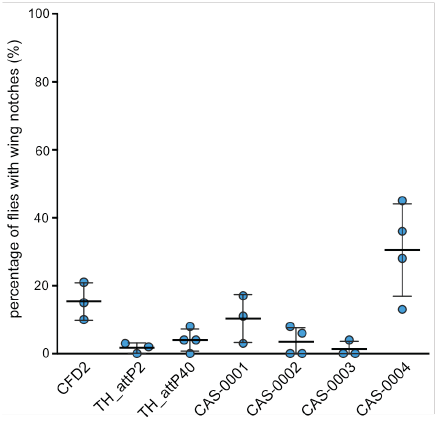
Assessing somatic activity of *nos-cas9* lines in combination with *U6:3-gRNA-wls*. Flies expressing the indicated *nos-cas9* line and *U6:3-gRNA-wls* were examined and the number of flies with notches of the wing margin counted. Dots represent data from individual crosses with mean ± SD indicated as lines. Notches were always small (< 10% of wing area) and only present in the minority of animals, demonstrating low levels of somatic activity. Typically, 30 - 50 flies were examined for each cross.

**Supplementary Figure 2 :**
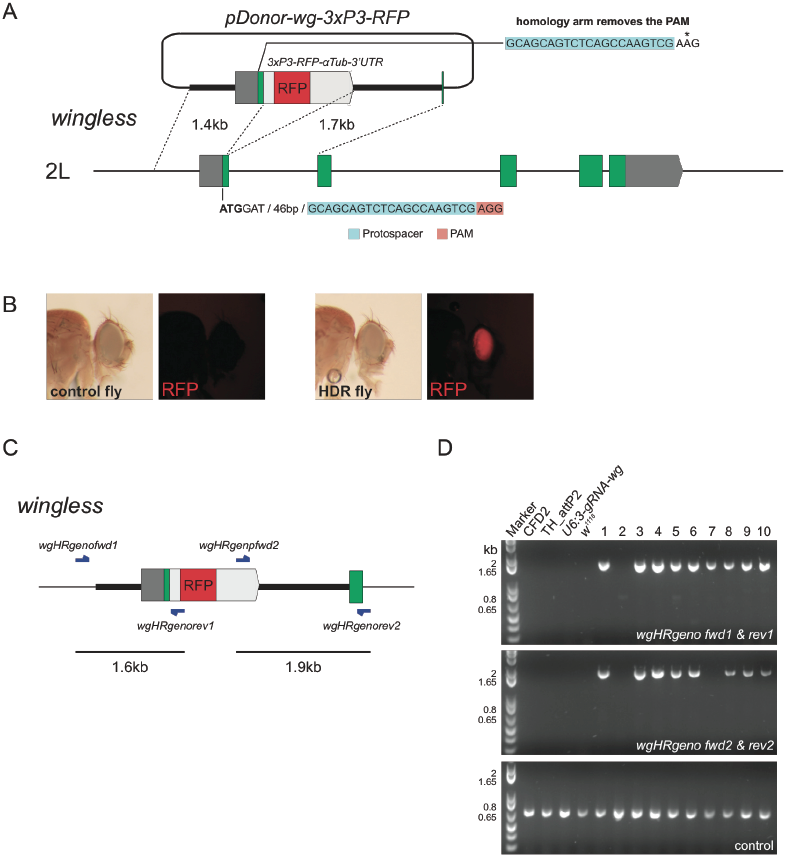
Targeted donor integration by HDR. (**A**) Schematic of the RFP donor vector in relation to the *wg* locus on chromosome arm 2L. The donor plasmid is based on a previous plasmid (Port et al., 2014), which was used to create a functional GFP knock-in in the *wg* locus. In the current plasmid, GFP was replaced by an autonomous RFP expression cassette. Integration of the donor plasmid causes a disruption of the *wg* gene and removes the protospacer adjacent motif (PAM) that is essential for cleavage by *gRNA-wg*, thus preventing recutting of the targeted allele. (**B**) Flies with genomic insertions of the donor plasmid can be easily identified by strong red fluorescence in the eye compared to control animals. (**C**) Schematic showing the *wg* locus after successful ends-out integration of the donor sequence. The indicated primers were used to analyse whether donor integration in RFP positive flies occurred at the correct location. Note that primers *wgHRgenofwd1* and *wgHRgenorev2* anneal outside the homology arms that are present in the donor plasmid. (**D**) A representative example of diagnostic PCRs from 10 flies from experiments designed to integrate the donor plasmid at the *wg* locus. Eight out of 10 flies gave rise to a product of the expected size with both *wgHRgeno* primer pairs and were scored as positive for RFP integration in *wg*. Only one PCR product was amplified from fly number 7, suggesting a complex integration event. Genomic DNA from control flies did not result in any PCR product with primers diagnostic for donor insertion in the *wg* locus (four lanes to the left of 1 – 10). Amplification of part of the *wls* locus was used as a positive control for DNA quality (bottom panel). A summary of all HDR genotyping experiments can be found in Suppl. Table 2.

**Supplementary Figure 3:**
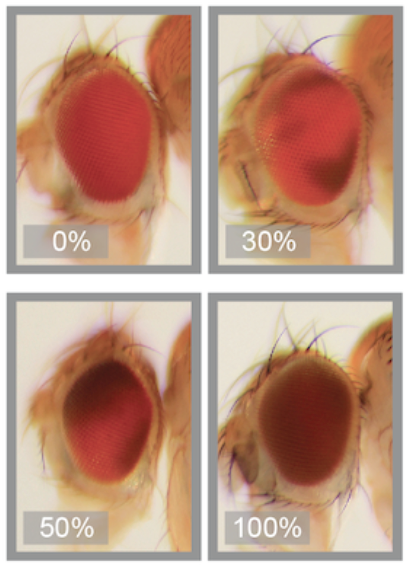
Assessing activity of gRNAs targeting *se*. Eyes from flies heterozygous for *act-cas9* and *gRNA-se* transgenes were examined by three independent researchers. Example images are shown of eyes with various degree of *se* mutant tissue, which has darker coloration than wild type tissue. Mean values from the independent observations are shown in Figure 3 (see also Suppl. Table 3).

**Supplementary Figure 4:**
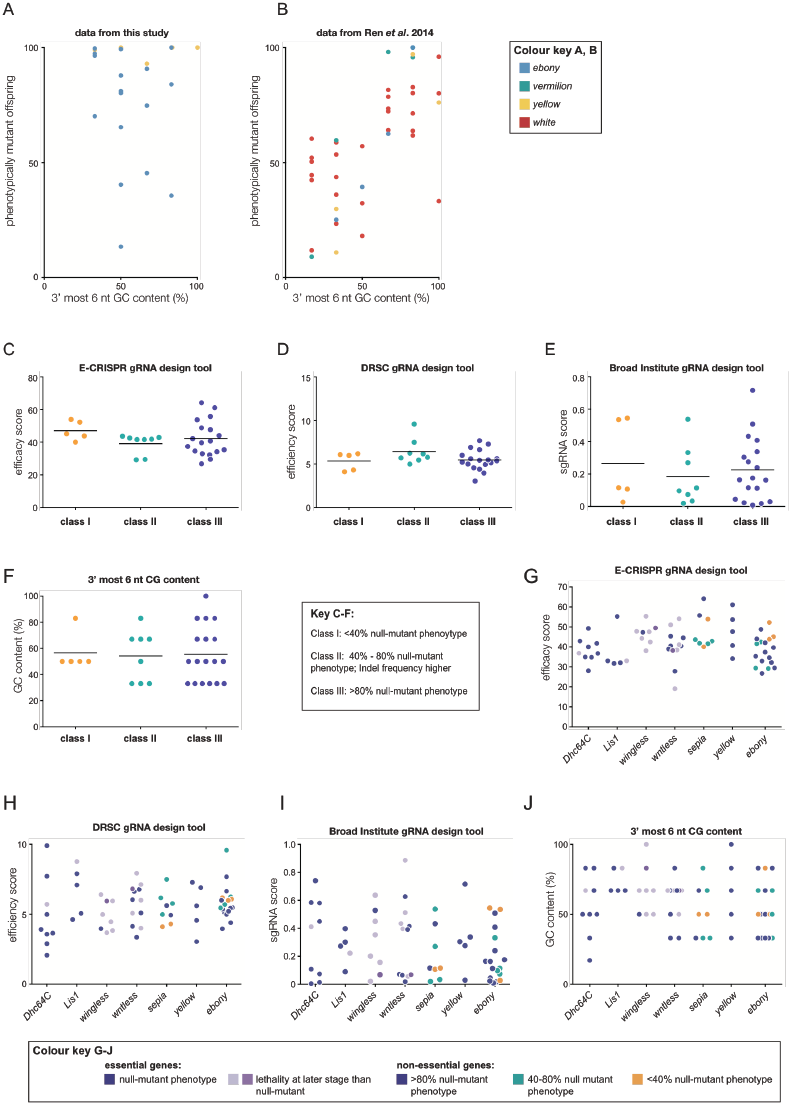
gRNA activity in CRISPR-it is not reliably predicted by online design tools or PAM proximal GC content of the target site. (**A** - **B**) Comparison of data from transgenic gRNAs from this study (A) and injected gRNAs from Ren *et al*. 2014 (B) relating the PAM proximal GC content of the target site (3’ most 6 nt of the spacer) to the rate of germ line transmission of non-functional alleles in phenotypic assays. Note that transmission of non-functional alleles was plotted as Ren *et al*. did not investigate in-frame functional mutations in their study. *act-cas9* was used in our study and *nos-cas9* (TH_attP2) was used by Ren *et al*.. These *cas9* sources had very similar activities when compared with the same gRNA transgene (Figure 1). The mean (± SD) rate of germ line transmission of non-functional alleles in non-essential genes was 80% ± 25 (N = 22) for our study and 57% ± 26 (N = 39) for Ren *et al*. (P = 0.0014, two-tailed unpaired t-test). (**C** – **F**) Prediction of gRNA activity with popular design tools. Activity scores for gRNA targeting *se*, *y* and *e* were generated with each online gRNA design tool (C – E; see Suppl. Table 4 for URLs); the GC content of the 3’ most 6 nt of the gRNA spacer was calculated manually (F). Values were plotted for gRNAs assigned to three classes (colour coded as in Figure 3). Whereas class I gRNAs mediate suboptimal mutagenesis, class II and III gRNAs mediate near maximal mutagenesis, but differ in phenotypic penetrance due to in-frame mutations. None of the methods can reliably predict reduced activity of the class I gRNAs. (G – J) gRNA activity scores plotted for individual pigmentation genes shows that lack of correlation in C – F is not caused by a single target gene. The four evaluated prediction methods for gRNA activity also failed to discriminate between those gRNAs targeting essential genes that gave a completely penetrant phenotype and those that did not. It should be noted, however, that the contribution of functional in-frame mutations to incompletely penetrant phenotypes is not known for these gRNAs.

**Supplementary Table 1:** Transgenic *cas9* lines used in this study.

| Construct(s) | Name (s) | Cas9 coding sequence<sup>a</sup><br>(Addgene plasmid number) | Integration site of cas9 construct<br>(chromosome arm) | Reference |
| --- | --- | --- | --- | --- |
| *act-cas9\** | CFD1; BL54590 | Hs_Cas9 (41815, 62209) | attP-ZH2a (X) | Port et al. (2014) |
| *vasa-cas9* | BL56552 | Hs_Cas9 (42230) | attP-VK00037 (2L) | Gratz et al. (2014) |
| *vasa-cas9\** | BL51324 | Hs_Cas9 (42230) | attP-VK00027 (3R) | Gratz et al. (2014) |
| *vasa-cas9\** | BL51323 | Hs_Cas9 (42230) | attP-ZH2a (X) | Gratz et al. (2014) |
| *vasa-cas9* | BL52669 | Hs_Cas9 (41815) | attP-ZH2a (X) | Sebo et al. (2014) |
| *nos-cas9\** | CFD2; BL54591 | Hs_Cas9 (41815, 62208) | attP-ZH2a (X) | Port et al. (2014) |
| *nosG4VP16 UAS-cas9\** | CFD3_nos; BL54593 | Dm-Cas9 | attP2 (3L) | Port et al. (2014) |
| *nos-cas9:GFP\** | CFD8 | Hs_Cas9:GFP (42234) | attP-ZH2a (X) | Port et al. (2014) |
| *nos-cas9* | TH00787.N | Dr_Cas9 | attP2 (3L) | Ren et al. (2013) |
| *nos-cas9* | TH00788.N | Dr_Cas9 | attP40 (2L) | Ren et al. (2013) |
| *nos-cas9* | CAS-0001 | Hs_Cas9 (41815) | attP40 (2L) | Kondo and Ueda (2013) |
| *nos-cas9* | CAS-0002 | Hs_Cas9 (41815) | Random insertion (X) | Kondo and Ueda (2013) |
| *nos-cas9* | CAS-0003 | Hs_Cas9 (41815) | Random insertion (3) | Kondo and Ueda (2013) |
| *nos-cas9* | CAS-0004 | Hs_Cas9 (41815) | Random insertion (2<br>(CyO balancer)) | Kondo and Ueda (2013) |

Abbreviations: *act* – *actin5C*; *nos* – *nanos*; Hs – *Homo sapiens*; Dm – *Drosophila melanogaster*; Dr – *Danio rerio*; BL – Bloomington *Drosophila* Stock Center stock number.
a, species refers to codon optimisation; all constructs express *Streptococcus pyogenes* Cas9 protein. \*, lines that were also evaluated in Port *et al.* 2014.

**Supplementary Table 2:** Summary of HDR results.

| Genotype of injected embryos | Injected plasmid(s) | DNA concentration gRNA/donor (ng/μl) | Independent experiments | No. of injected embryos | Embryo survival rate (%) | No. of G<sub>0</sub> males crossed | No. of sterile or dead G<sub>0</sub> males | No. of male founders/fertile G<sub>0</sub> males screened (%) | No. of RFP-positive F<sub>1</sub> males/total screened (%) | No. of confirmed knock-ins into *wg*/total screened (%)<sup>a</sup> |
| --- | --- | --- | --- | --- | --- | --- | --- | --- | --- | --- |
| *nos-cas9* CFD2, *U6:3-gRNA-wg* | *pDonor wg-3P3-RFP* | 500 | Expt 1 | 125 | nd | 10 | 2 | 7/8 (88) | 41/180 (23) | 35/36 (97) |
| *nos-cas9* CFD2, *U6:3-gRNA-wg* | *pDonor wg-3P3-RFP* | 100 | Expt 1 | 151 | 30 | 13 | 0 | 9/13 (69) | 46/242 (19) | 7/8 (88) |
| *nos-cas9* TH_attP2, *U6:3-gRNA-wg* | *pDonor wg-3P3-RFP* | 100 | Expt 1 | 131 | 18 | 9 | 2 | 7/7 (100) | 43/123 (35) | 7/8 (88) |
|  |  |  | Expt 2 | 149 | 32 | 10 | 1 | 5/9 (56) | 13/97 (13) | 5/6 (83) |
|  |  |  | **Total** | **280** | **25** | **19** | **3** | **12/16 (75)** | **56/220 (25)** | **12/14 (86)** |
| *nos-cas9* CFD2 | *pU6:3-gRNA-wg/*<br>*pDonor wg-3P3-RFP* | 500/500 | Expt 1 | 91 | nd | 10 | 2 | 7/8 (88) | 19/222 (9) | 14/18 (78) |
| *nos-cas9* CFD2 | *pU6:3-gRNA-wg/*<br>*pDonor wg-3P3-RFP* | 75/100 | Expt 1 | 136 | 32 | 16 | 0 | 8/16 (50) | 18/232 (8) | 5/6 (83) |
|  |  |  | Expt 2 | 117 | 34 | 8 | 0 | 3/8 (38) | 9/270 (3) | 4/6 (67) |
|  |  |  | **Total** | **253** | **33** | **24** | **0** | **11/24 (46)** | **27/502 (5)** | **9/12 (75)** |
| *nos-cas9* TH_attP2 | *pU6:3-gRNA-wg/*<br>*pDonor wg-3P3-RFP* | 75/100 | Expt 1 | 134 | 22 | 11 | 3 | 6/8 (75) | 33/217 (15) | 4/4 (100) |
| *w*<sup>1118</sup> | *pDCC6-gRNA-wg/*<br>*pDonor wg-3P3-RFP* | 75/100 | Expt 1 | 268 | 7 | 3 | 1 | 2/2 (100) | 6/46 (13) | 3/3 (100) |
|  |  |  | Expt 2 | 123 | 52 | 26 | 5 | 2/21 (10) | 6/322 (2) | 3/3 (100) |
|  |  |  | **Total** | **391** | **22** | **29** | **6** | **4/23 (17)** | **12/368 (3)** | **6/6 (100)** |

Abbreviations: nd – not determined; Expt – experiment.

**Supplementary Table 3:**
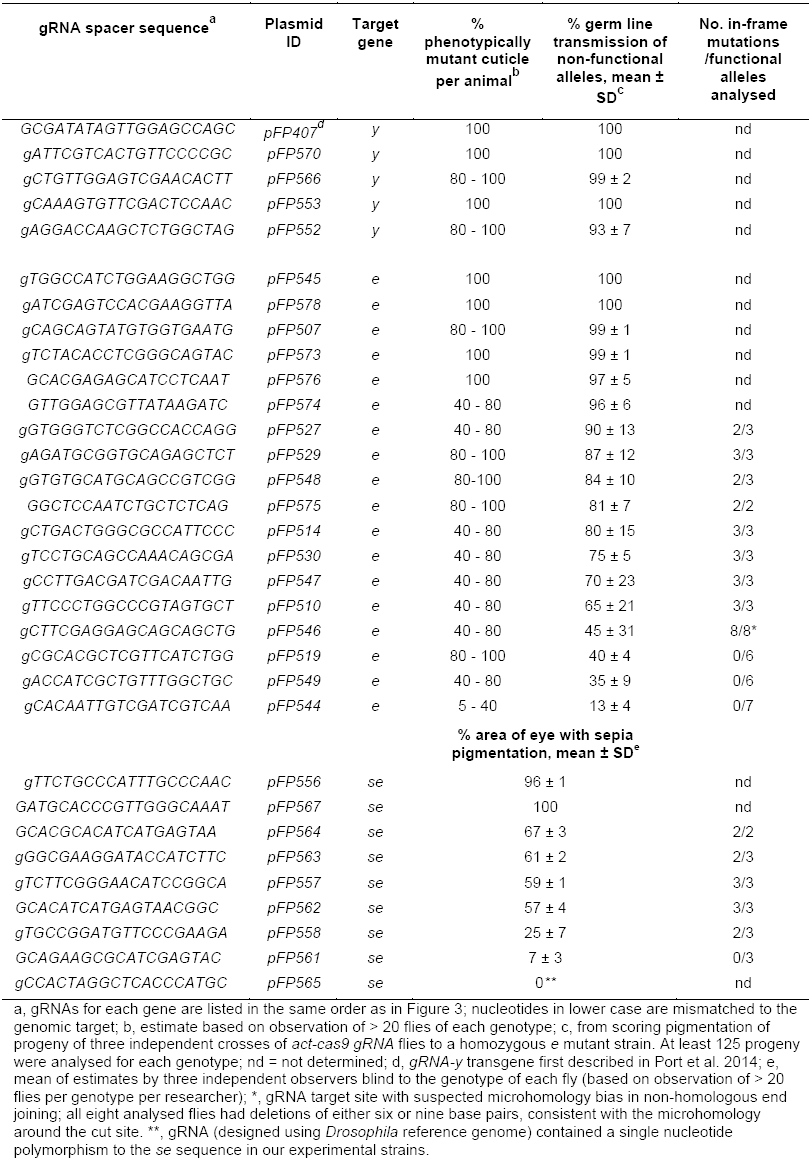
Assessment of somatic and germ line mutagenesis mediated by *U6:3-gRNA* transgenes targeting pigmentation genes.

**Supplementary Table 4:**
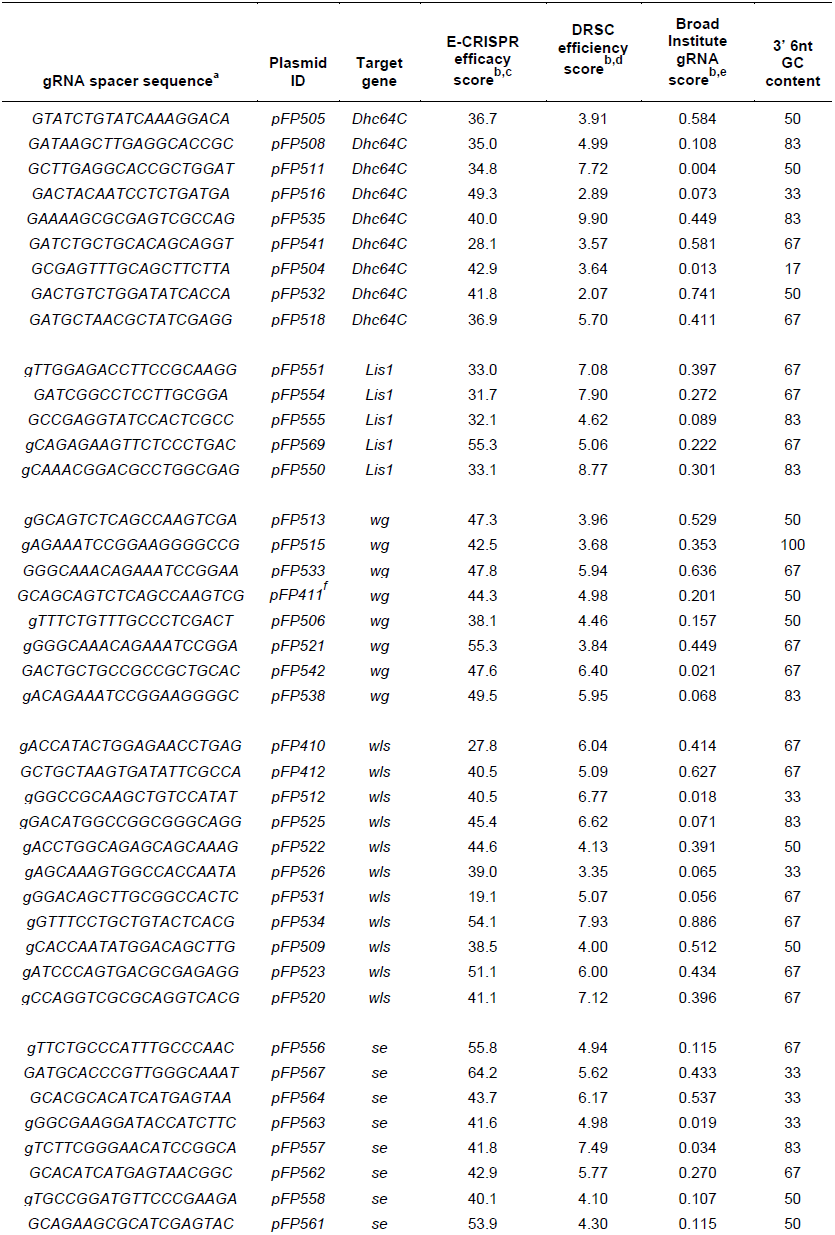
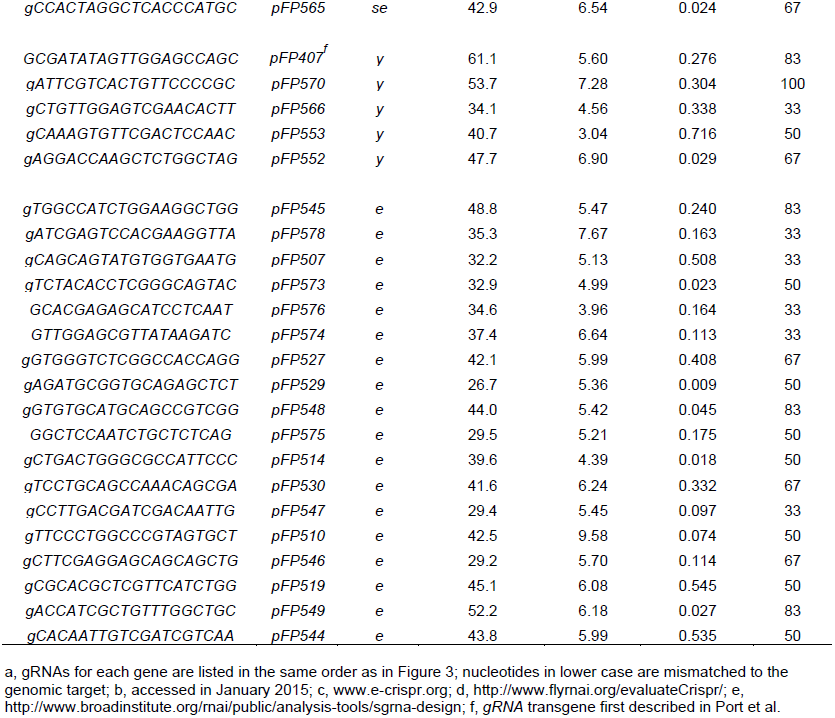
Computed gRNA efficiency scores and 3’ 6nt GC content.

**Supplementary Table 5:**
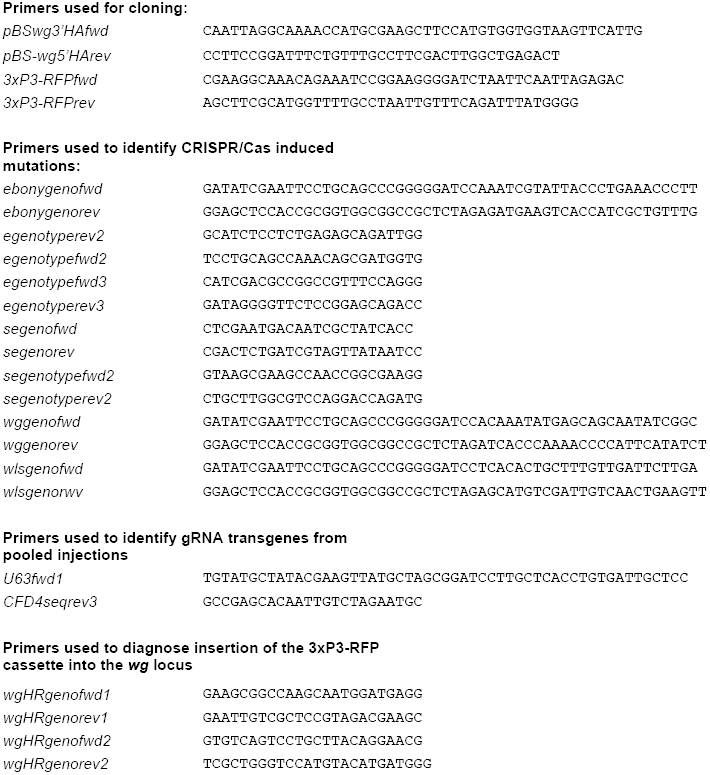
Oligonucleotides used in this study (5’ – 3’)

